# The landscape of antibody binding affinity in SARS-CoV-2 Omicron BA.1 evolution

**DOI:** 10.1101/2022.09.13.507781

**Authors:** Alief Moulana, Thomas Dupic, Angela M. Phillips, Jeffrey Chang, Anne A. Roffler, Allison J. Greaney, Tyler N. Starr, Jesse D. Bloom, Michael M. Desai

## Abstract

The Omicron BA.1 variant of SARS-CoV-2 escapes convalescent sera and monoclonal antibodies that are effective against earlier strains of the virus. This immune evasion is largely a consequence of mutations in the BA.1 receptor binding domain (RBD), the major antigenic target of SARS-CoV-2. Previous studies have identified several key RBD mutations leading to escape from most antibodies. However, little is known about how these escape mutations interact with each other and with other mutations in the RBD. Here, we systematically map these interactions by measuring the binding affinity of all possible combinations of these 15 RBD mutations (2^15^ = 32,768 genotypes) to four monoclonal antibodies (LY-CoV016, LY-CoV555, REGN10987, and S309) with distinct epitopes. We find that BA.1 can lose affinity to diverse antibodies by acquiring a few large-effect mutations and can reduce affinity to others through several small-effect mutations. However, our results also reveal alternative pathways to antibody escape that do not include every large-effect mutation. Moreover, epistatic interactions are shown to constrain affinity decline in S309 but only modestly shape the affinity landscapes of other antibodies. Together with previous work on the ACE2 affinity landscape, our results suggest that escape of each antibody is mediated by distinct groups of mutations, whose deleterious effects on ACE2 affinity are compensated by another distinct group of mutations (most notably Q498R and N501Y).

---

In November 2021, the SARS-CoV-2 Omicron BA.1 variant emerged and quickly rose to high frequency worldwide, in part due to its ability to escape preexisting immunity^1–4^. This immune escape is mediated by mutations in the receptor binding domain (RBD) of the spike protein, which is the major target of SARS-CoV-2 neutralizing antibodies^5–8^. Antibodies targeting the RBD can bind different epitopes, and they have been grouped into several classes^9,10^. Some previous SARS-CoV-2 variants which have a subset of the fifteen mutations found in the BA.1 RBD (e.g. K417N, N501Y in Beta and T478K, N501Y in Delta) can evade some antibodies of certain epitope classes but still bind to others^11–14^. In contrast, BA.1 can escape most antibodies that bind to very distinct epitopes, including antibodies elicited by previously circulating variants^1,15,16^.

Existing studies of SARS-CoV-2 immune escape have focused on measuring the effects of single mutations (or, in some cases, of a small subset of mutations) on antibody escape in the context of specific SARS-CoV-2 variants^15,17,18^. However, simultaneous escape of most antibodies is likely to require multiple mutations, and it is unclear how these mutations might interact. A large body of work has demonstrated that the specific combination of mutations in the BA.1 variant can evade various antibodies of distinct epitopes^1,3,15,19^. However, the landscape on which this evolution occurred is not well understood. Do mutations involved in escape from one antibody with a certain epitope interfere with the effects of those involved in escaping others with different contact sites, or are the effects largely independent? And how are these effects mediated by epistatic interactions with other mutations in the RBD?

As we observed in previous work, several of these antibody-escape mutations also reduce affinity to ACE2, suggesting that they were positively selected because they contribute to immune escape^20–22^. Importantly, epistatic interactions between these mutations dramatically impact ACE2 affinity and may also differentially impact the escape of antibodies with very different epitopes^23,24^. For example, escape from some antibodies like S309 has been difficult to attribute to specific mutations^22,25^, perhaps because measurements have so far been limited to single mutations. These observations suggest that we need to more comprehensively characterize the role of epistasis and potential trade-offs to understand the simultaneous evolution of escape from multiple antibodies of distinct epitopes and ACE2 binding affinity.

Here, to understand how immune pressure may have shaped the evolution of BA.1, we measured the equilibrium binding affinities (*K*_D, app_) of the spike protein RBD to four therapeutic monoclonal antibodies (mAbs) with distinct RBD epitopes: LY-CoV016, LY-CoV555, REGN10987, and S309, for all possible evolutionary intermediates between the ancestral Wuhan Hu-1 RBD and the BA.1 variant. This set of antibodies includes the primary epitopes generally covered by therapeutic mAbs^10,16^. The first three antibodies are fully escaped by Omicron BA.1, while S309 has reduced affinity. We find that for each antibody, only a few mutations significantly impact affinity, and these mutations are largely (but not entirely) orthogonal between the four antibodies. Additionally, we find that epistasis plays a limited role in determining affinity to antibodies that are fully escaped by BA.1 but contributes substantially to the reduced affinity for the partially escaped antibody, S309. Together, this work systematically characterizes how SARS-CoV-2 can evade distinct RBD-targeted antibodies while maintaining ACE2 affinity.

## RESULTS

In previous work, we generated a combinatorically complete library comprising all possible intermediates between the ancestral SARS-CoV-2 Wuhan Hu-1 spike protein RBD and the Omicron BA.1 variant^23^. The BA.1 RBD differs from Wuhan-1 by fifteen amino acid substitutions, so this library contains 2^15^ variants containing all possible combinations of these fifteen mutations. This RBD library is displayed on the surface of yeast, such that each yeast cell expresses a single variant. Here, we use Tite-Seq (a high-throughput method that integrates flow cytometry and sequencing^23,26–28^; see Supplementary Figure S1A) to measure the equilibrium binding affinities of all 32,768 variants to four antibodies with different epitopes (LY-CoV016, LY-CoV555, REGN10987, and S309). The resulting *K*_D, app_ correlates between biological duplicates and with isogenic measurements made by flow cytometry (Supplementary Figure S1A, Supplementary Table S1).

Of the 32,768 variants in our library, we obtain *K*_D, app_ for at least ~30,000 variants to each of the mAbs (32,112 for Ly-CoV016, 30479 for REGN10987, 29892 for CoV555, and 32602 for S309) after removing variants with poor titration curves (r^2^ < 0.8 or σ > 1; see Methods). These *K*_D,app_ range from 0.1 nM to 1 μM (which is our limit of detection and likely corresponds to nonspecific binding), with 50% of the variants fully escaping LY-CoV016 (defined as having *K*_D,app_ above the limit of detection), 55% fully escaping LY-CoV555, 33% fully escaping REGN10897, and no variants fully escaping S309 (Figure 1A; see https://desai-lab.github.io/wuhan_to_omicron/ for an interactive data browser). Escape from LY-CoV016, LY-CoV555, and REGN10897 is mediated by one or a few strong-effect mutations, with other mutations more subtly impacting affinity (Figure 1B). In general, strong-effect mutations make substantial contact with the corresponding antibody. Consistent with previous studies^14,16,17,29^, these strong-effect mutations are largely distinct for each antibody, which presumably reflects their non-overlapping footprints on the RBD (Figure 1C) and suggests that evolution of escape from each antibody can be, to some extent, orthogonal.

**Figure 1.**
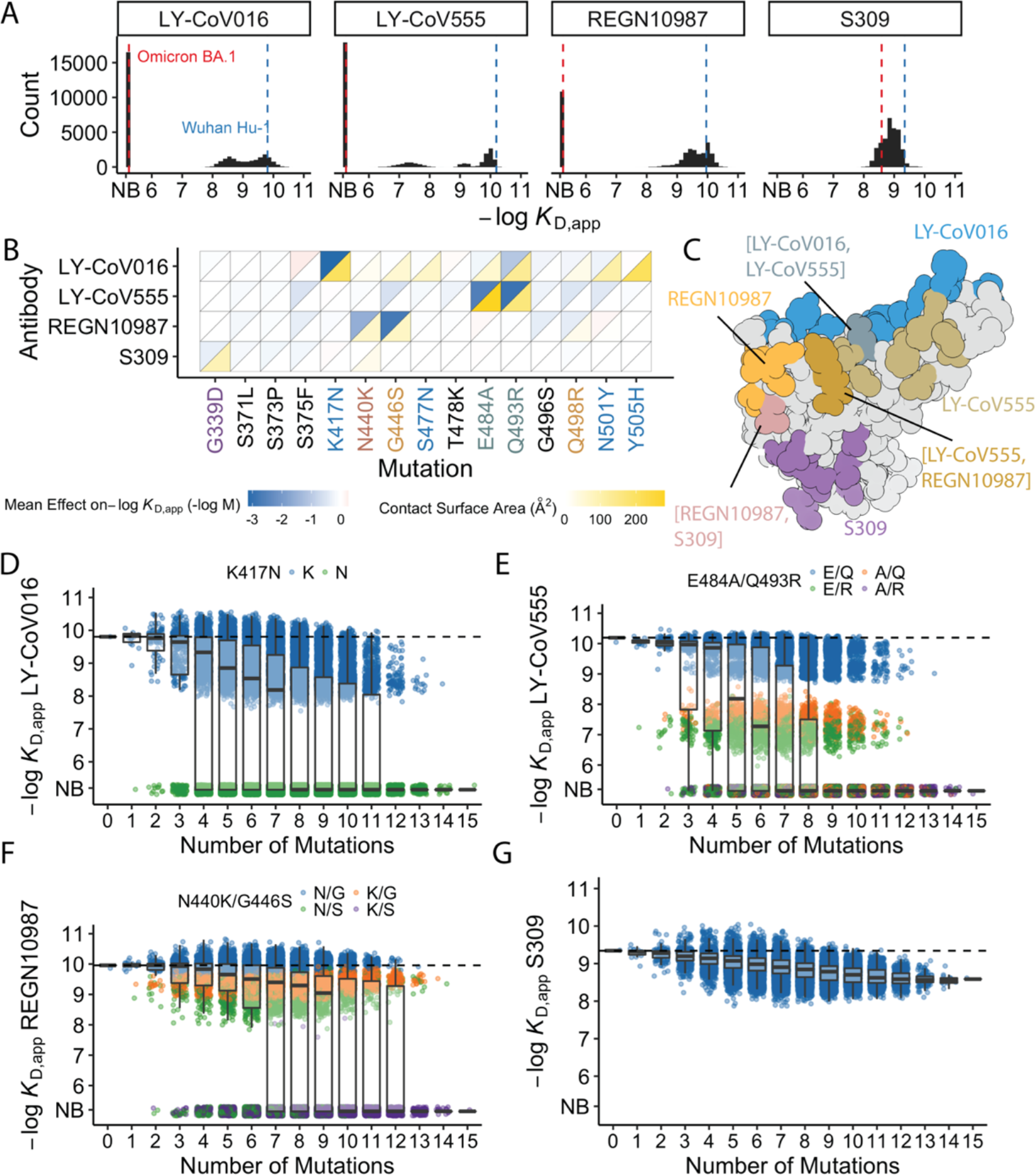
Antibody affinity landscape. (**A**) Binding affinities to one antibody from each class (LY-CoV016, LY-CoV555, REGN10987, and S309, from classes 1-4, respectively) across all N=32,768 RBD genotypes tested. Binding affinities are shown as −log*K*_D,app_; vertical blue and red dashed lines indicate the −log*K*_D,app_ for Wuhan Hu-1 and Omicron BA.1, respectively. ‘NB’ denotes non-binding (escape). (**B**) Mean effect of each mutation on antibody and ACE2 affinity (defined as the change in −log*K*_D,app_ resulting from mutation) plotted with contact surface area between each residue and each antibody. Mutations are colored by footprint highlighted in (**C**). (**C**) Structure of SARS-CoV-2 BA.1 RBD with each antibody footprint annotated (PDB ID 7KMG, 6WPT, 7C01, 6XDG). Residues with overlapping footprints are colored and labeled accordingly. (**D-G**) Distribution of binding affinities to different antibodies grouped by number of Omicron BA.1 mutations. Binding affinity of the Wuhan Hu-1 variant is indicated by horizontal dashed lines. Variants with antibody escape mutations of interest are colored as noted in each key. NB denotes non-binding (escape).

The picture is more complex for S309, where BA.1 has reduced affinity relative to Wuhan Hu-1, but ~19% of variants have lower affinity than BA.1. These differences are not attributable to one or two strong effect mutations (Figure 1A-B). In addition, although most mutations reduce affinity, three mutations have small positive effects (on average across all backgrounds at the other loci): S375F for LY-CoV016, E484A and N501Y for REGN10987 (Figure 1B). Intriguingly, each of these mutations reduces affinity to at least one of the other antibodies, and N501Y significantly improves binding to ACE2, suggesting a potential role for trade-offs (and/or epistasis that mitigates these effects on specific backgrounds).

For each antibody, binding affinities generally decrease as the number of mutations increase (Figure 1D-G). For LY-CoV016, LY-CoV555, and REGN10897 this trend is observed amongst variants with and without the large-effect escape mutations (Figure 1D-F). For LY-CoV016, K417N is sufficient for escape (Figure 1D, green), whereas both LY-CoV555 and REGN10987 require at least two mutations for complete escape. For LY-CoV555, both E484A and Q493R decrease affinity drastically (1000 and 100-fold, respectively), but only the combination of both mutations lead to complete escape. Complete escape from REGN10987 also requires two mutations (N440K and G446S) but the individual effects of these mutations are more subtle (reducing affinity by 10 and 5-fold, respectively). For S309, affinity declines after a few mutations and in some backgrounds increases upon further mutation, suggesting that interactions between these mutations are important in determining affinity (Figure 1G).

### Mostly orthogonal large-effect mutations

We first focused on analyzing how mutations and combinations of mutations lead to complete escape (defined as *K*_D,app_ above our limit of detection) for specific sets of antibodies. To do so, we analyze the enrichment of specific mutations among non-binders (Figure 2A). We find a largely orthogonal set of one or two mutations are enriched among variants that do not bind each antibody: almost all variants that do not bind LY-CoV016 contain K417N, almost all variants that do not bind REGN10987 contain G446S and many also contain N440K, and E484A and Q493R are highly enriched among variants that do not bind LY-CoV555. This suggests that the RBD can evolve to independently escape antibodies with each distinct epitope and mutations can to some extent act independently on binding to each antibody.

**Figure 2.**
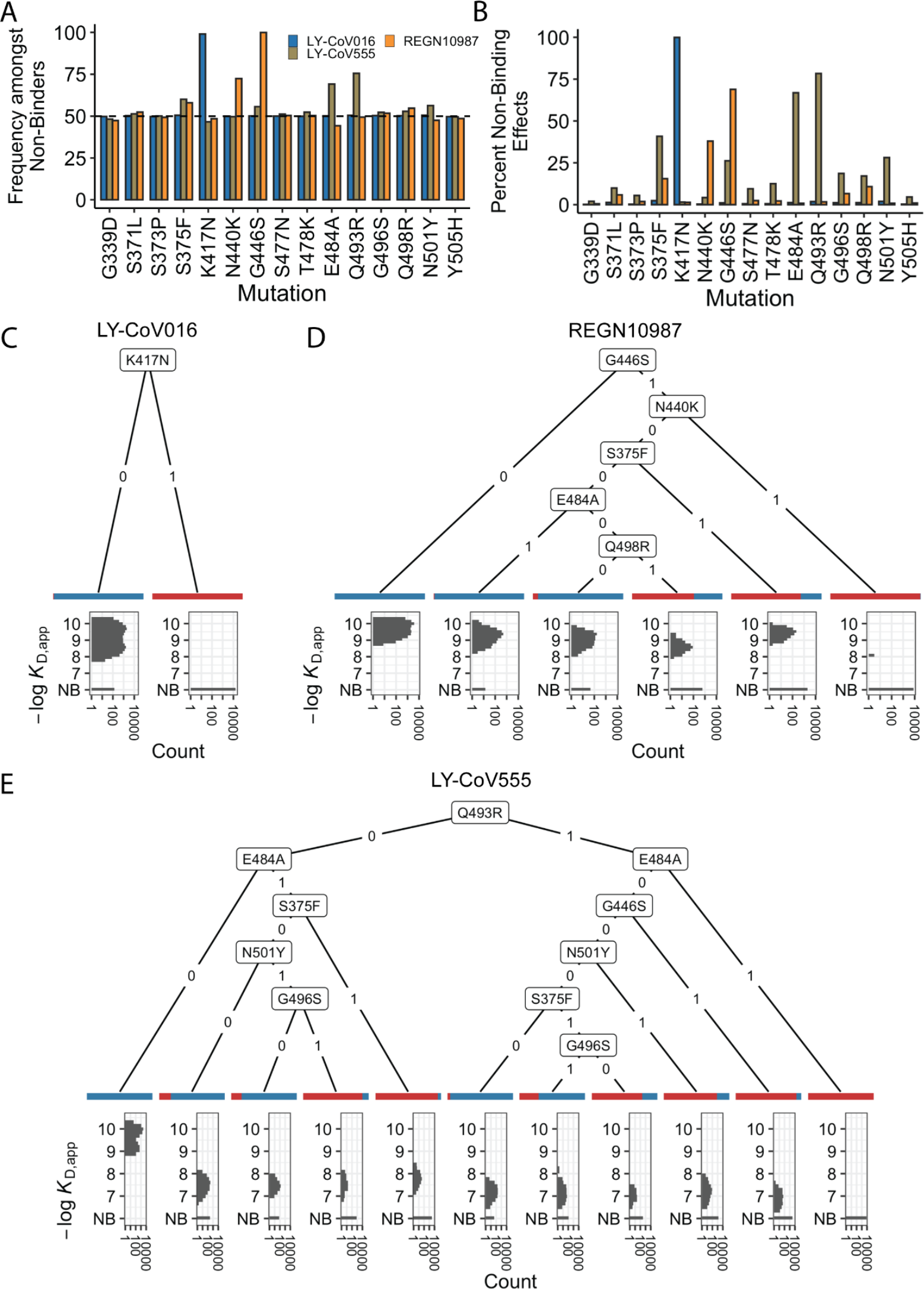
Escape mutations and genotypes. (**A**) Fraction of antibody-escaping genotypes with each mutation **(B)** Fraction of variants for which a given mutation confers antibody escape. Effects are colored as in (A). (**C-E**) Decision trees of escape phenotype for each antibody modeled as a function of the mutations present. Each leaf is annotated by the proportion of the genotypes that escape the corresponding antibody (red: escape, blue: does not) and by corresponding affinity distribution. NB denotes non-binding.

To analyze this further, we calculate the percentage of genetic backgrounds on which each mutation leads to complete escape from a specific antibody (i.e. that mutation converts a variant with measurable *K*_D,app_ to a *K*_D,app_ above our limit of detection). We see that for each antibody, one or two mutations abrogate binding. These sets of mutations are largely orthogonal among antibodies (Figure 2B), consistent with the enrichment analysis (Figure 2A). Specifically, K417N always abrogates binding to LY-CoV016, G446S and N440K often abrogate binding to REGN10987, and E484A and Q493R often abrogate binding to LY-CoV555.

However, we note that this orthogonality is not complete. For example, G446S is slightly enriched among LY-CoV555 binders, while mutation E484A is slightly depleted among variants that do not bind REGN10987 (Figure 2A). Consistent with this, G446S sometimes abrogates binding to LY-CoV555 (Figure 2B). In addition, some apparently smaller-effect mutations can be involved in abolishing binding to multiple antibodies. For example, S375F is weakly enriched among variants that do not bind REGN10987 and LY-CoV555 and often abrogates binding to these two antibodies, with G496S, Q498R, and N501Y also playing a role.

To summarize how these different mutations can act individually or in combination to lead to antibody escape, we inferred a decision tree to classify variants as binders or non-binders. To do so, for each antibody we calculate the mutation that maximally partitions the variants into binders or non-binders. If this partitioning is not perfect, we then calculate the second mutation that maximally partitions the variants conditional on each possible state of the first site. We then proceed to further partition variants based on additional mutations in the same way, until the variants are perfectly partitioned or no further mutations can significantly improve the partitioning (see Methods). We show the corresponding decision trees for LY-CoV016, REGN10987, and LY-CoV555 in Figures 2C, D, and E respectively. As expected, the tree associated with LY-CoV016 is very simple: the mutation K417N perfectly partitions the variants into binders and non-binders. In contrast, the trees for REGN10987 and LY-CoV555 have more complex structures, reflecting the fact that it is possible to abrogate affinity to these antibodies via multiple distinct combinations of mutations. For example, variants can escape REGN10987 by acquiring G446S and N440K (100%), or alternatively, with S375F and G446S (77%, as additional mutations are also required). For LY-CoV555, different sets of mutations can lead to escape (e.g. Q493R and G446S or E484A and S375F), and G496S can help or hinder escape depending on the presence of other mutations. Some of the mutations resulting in LY-CoV555 escape partially overlap with those for REGN10987 (i.e. they are not fully orthogonal), suggesting that selection pressure from one antibody could promote subsequent escape of another.

### Inference of epistatic affinity landscapes

In addition to large-effect mutations which lead to complete escape of specific antibodies, a variety of other sites contribute to more subtle but potentially important changes in binding affinities. To analyze these subtle effects as well as the large-effect mutations leading to escape, we defined a linear model for −log(*K*_D, app_) as the sum of single (additive) mutational effects plus interaction terms up to a specified order (note that because −log(*K*_D, app_) is proportional to the free energy of binding, we expect it to behave additively in the absence of epistatic interactions). Because non-binding variants have −log(*K*_D, app_) beyond our limit of detection, we fit a Tobit model (a class of regression model capable of handling truncated measurements, see Methods for details) using maximum likelihood with an L2-norm Lasso regularization. Specifically, we partition our data into training (90%) and test (10%) sets and use the training dataset to fit epistatic coefficients to a linear model truncated at each order (e.g. truncating to first-order yields additive mutational effects, second-order includes both additive effects and pairwise terms, and so on). We then evaluate performance (as the coefficient of variation) of each model on the held-out test dataset, and compare the model performance using −log(*K*_D, app_) for each of the antibodies and ACE2 (Figure 3A).

**Figure 3.**
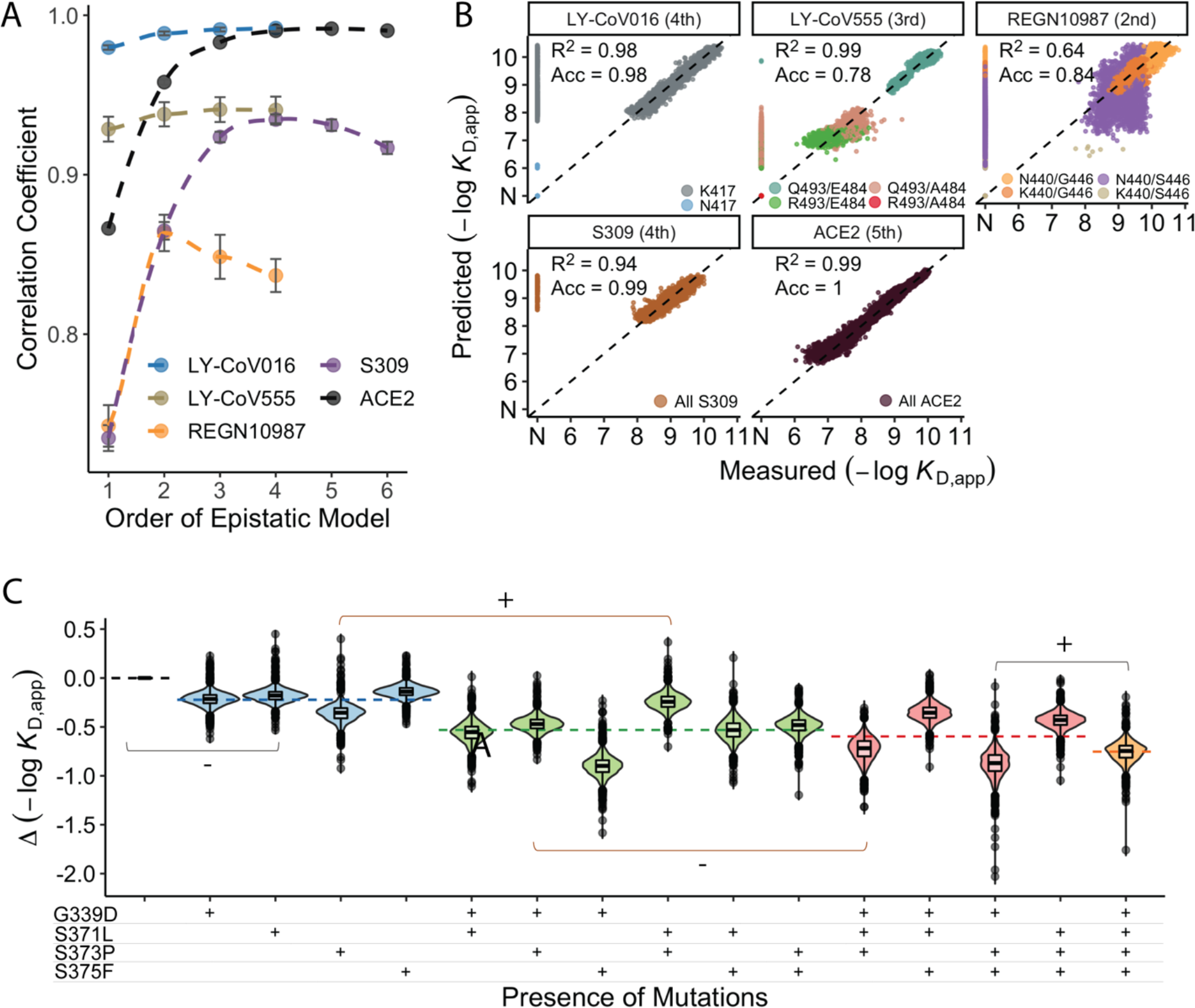
Epistastic effects on antibody binding. (**A**) Correlation coefficients between the measured values of −log(*K*_D,app_) and the model estimate for various orders of epistatic models. Correlations are computed on the hold-out subset averaged over 10 folds of cross-validation. zoomed-in version for orders 3 to 6. (**B**) Binding affinities predicted by complete coefficients from the optimum epistasis model are compared to the measured binding affinities for each antibody. Points are colored by mutations present in the genotypes, ‘N’ corresponds to nonbinding genotypes. The accuracy measures the quality of the binary classification between binders and non-binders and the coefficient of determination R^2^ refer to the correlation between inferred and measured binding affinities, excluding non-binders. (**C**) Effects of mutations G339D, S371L, S373P, and S375F on S309 affinity grouped by the presence of each mutation. Each violin color corresponds to the number of mutations considered. Dashed line color denotes the average effect for each group represented by the violin color.

We find that adding epistatic interactions improves the predictive power of the model for all four antibodies as well as for binding to ACE2, though the optimal order varies (Figure 3A). This indicates that epistasis does play a significant role in all cases (up to second order for REGN10987, to third order for LY-CoV555, and to fourth or higher order for LY-CoV016, S309, and ACE2). The additive, pairwise, and higher-order coefficients resulting from these models are summarized in Supplementary Figure S2. In general, we find many strong interactions across several positions in each antibody, involving both the sites that strongly determine escape variants for that antibody (e.g. between N440K and G446S for REGN10987) as well as others.

Notably, we find that the higher-order epistasis plays a much stronger role in determining affinity for ACE2 than for the three antibodies fully escaped by BA.1 (Figure 3B). This reflects the impact of a few strong-effect mutations in determining affinity for LY-CoV555, LY-CoV016, and REGN10987, and the role of compensatory epistasis in determining ACE2 affinity. In other words, while epistasis is relevant for all measured phenotypes, antibody escape is more simply determined by the additive effects of individual mutations, while maintaining ACE2 affinity involves more complex epistatic interactions.

High-order epistatic interactions are also important in determining affinity to S309. In Figure 3C we highlight four neighboring mutations (G339D, S371L, S373P, and S375F) which interact non-additively to produce the reduction in affinity observed in BA.1 relative to Wuhan Hu-1. Each of these mutations weakly reduces affinity on their own, and specific combinations of these mutations can reduce affinity by up to two orders of magnitude, but the reduction in affinity resulting from all four mutations is less than some sets of three mutations. These patterns emerge from a complex set of high-order epistatic interactions among the mutations. For example, S371L reduces affinity on the Wuhan Hu-1 background but increases affinity on the background containing G339D, S373P, and S375F (and without S375F, S371L increases affinity in the presence of S373P if G339D is absent but not if it is present). Thus, some variants lacking S371L evade S309 more effectively than BA.1, and interestingly we note that this mutation is absent in BA.2 and BA.3 and replaced instead by S371F.

### Tradeoffs between antibody and ACE2 affinities

In previous work, we found that antibody escape mutations (as defined in earlier studies) typically reduce ACE2 affinity, suggesting that viral evolution is constrained by a tradeoff between immune evasion and the ability to enter host cells. Consistent with this, we find here that variants that escape one or more antibodies (as defined by the data reported in this work) but have few additional mutations have reduced ACE2 affinity relative to Wuhan Hu-1. However, as additional BA.1 mutations are accumulated, the ACE2 binding affinity tends to increase until it exceeds the Wuhan Hu-1 value even in the presence of multiple antibody escape mutations (Figure 4A). This suggests that the evolution of the BA.1 variant is driven both by immune escape and the need for compensatory mutations that mitigate the negative effects of the escape mutations on ACE2 binding.

**Figure 4.**
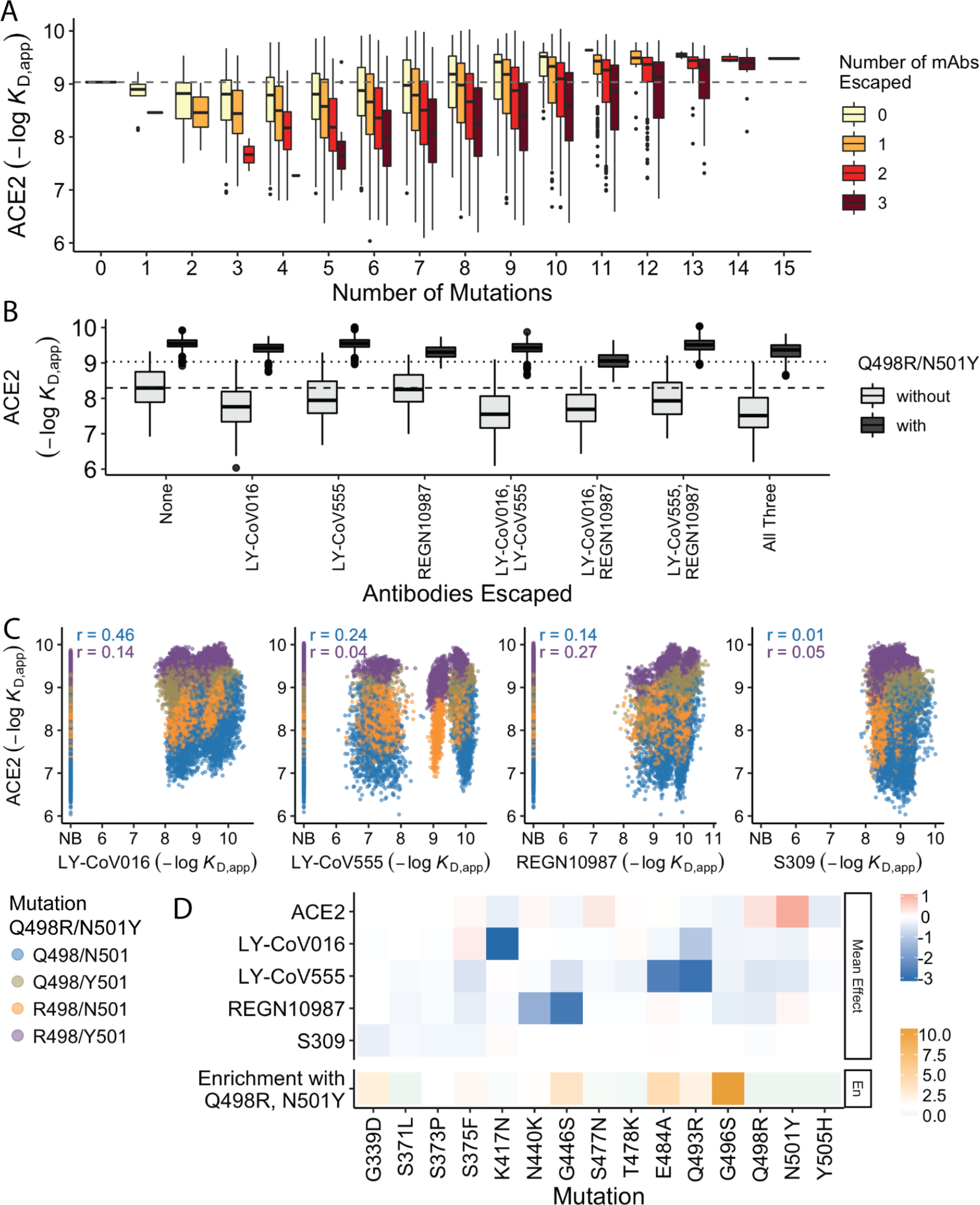
Trade-offs and comparison with ACE2 affinity. (**A**) Distribution of ACE2 binding affinity grouped by number of BA.1 mutations and the number of monoclonal antibodies escaped. (**B**) ACE2 affinity distribution grouped by antibodies escaped and the presence of compensatory mutations (N501Y and Q498R). (**C**) Affinities to monoclonal antibodies plotted as a function of the ACE2 affinity for all genotypes. Points are colored by presence of Q498R and N501Y. **(D)** Mean effect of each mutation on antibody affinity and on ACE2 affinity (red-blue colormap) compared to the enrichment of their frequency with Q498R and N501Y (orange colormap). The enrichment score is defined as the normalized frequency a mutation emerged on a branch on which mutations Q498R and N501Y appear divided by the normalized frequency it emerged on any intermediate background between Wuhan and BA.1.

The strength of this tradeoff and the potential importance of compensatory evolution is distinct between the different antibodies (Figure 4B,C). For example, escape from LY-CoV016 or LY-CoV555 reduces ACE2 binding affinity in the absence of compensatory mutations (Q498R and N501Y), but not in their presence (Figure 4B). In contrast, REGN10987 escape does not strongly reduce affinity to ACE2, whether or not Q498R and N501Y are present. However, this tradeoff is likely relevant overall, as escaping all three antibodies substantially reduces ACE2 affinity in the absence of Q498R and N501Y, while the reduction in ACE2 affinity is minimal in their presence. Consistent with this general picture, the frequency of most escape mutations (G446S, E484A, and Q493R) is higher across the SARS-CoV-2 phylogeny in the presence of compensatory mutations (Figure 4D), though we note that this is not universally true (e.g. the frequency of N440K is only slightly higher in the presence of compensatory mutations, and the frequency of K417N is lower with the compensatory mutations).

Although antibody escape mutations do tend to reduce ACE2 affinity, antibody binding affinity (but not complete escape) is not strongly correlated with ACE2 affinity (Figure 4C). The details of this relationship vary by antibody. For LY-CoV016 and LY–CoV555, there is a weak overall positive correlation (i.e. lower antibody affinity also tends to correspond to reduced ACE2 affinity). However, this correlation is dominated by the variants that lack the compensatory mutations at sites 498 and 501; in the presence of Q498R and N501Y the correlation largely disappears. For REGN10987, there is a similar weak overall positive correlation, which is less dependent on Q498R and N501Y. Finally, for S309, there is essentially no correlation regardless of whether compensatory mutations are present. In other words, for this antibody it appears possible to evolve reduced affinity (at least to the extent seen in BA.1) without compromising ACE2 binding. We also note that while compensatory mutations Q498R and N501Y largely drive the variance in ACE2 affinity, they minimally impact antibody binding affinities.

## DISCUSSION

Overall, we find that BA.1 escape from LY-CoV016, LY-CoV555, and REGN10987 is driven by a relatively small set of mutations: K417N for LY-CoV016, N440K and G446S for REGN10987, E484A and Q493R for LY-CoV555. These mutations have largely orthogonal effects on affinity to the three antibodies, suggesting that the evolution of escape to each can occur independently, as might be expected given the distinct epitopes they target^9,16,18^.

However, despite these largely orthogonal effects of large-effect mutations on antibody escape, we do observe limited trade-offs between LY-CoV016 and LY-CoV555 and between LY-CoV555 and REGN10987, with three mutations (S375F, K417N, and E484A) improving affinity to one antibody and reducing affinity to another. These positive effects are modest compared to the reductions in affinity caused by other mutations. In fact, for all antibodies studied here, outside of a few large-effect mutations that abrogate or nearly abrogate binding, most mutations weakly impact binding affinity and, even collectively, are insufficient to abrogate it.

In contrast to the orthogonality of antibody escape, trade-offs between binding ACE2 and escaping antibodies are much stronger. While the mutations with a small effect on antibody escape are mostly uncorrelated to ACE2 affinity, the strong-effect mutations substantially reduce ACE2 affinity. Thus, ACE2 affinity is lower for variants that escape a larger number of antibodies, unless compensatory mutations are acquired, suggesting that these compensatory mutations potentiated the establishment of the antibody escape mutations^23,30^.

We also find that epistatic interactions are important in determining antibody affinity. This is particularly true for S309. Prior to this work, the reduced affinity of S309 to Omicron could not be attributed to specific mutations^16,25^. Here, we find that this ambiguity can be resolved by examining higher-order interactions between mutations, as the reduction in affinity is attributable to a fourth-order epistatic interaction. This finding suggests that the potential for future SARS-CoV-2 lineages to escape S309 and similar antibodies could depend on epistatic interactions between emerging mutations.

We note that our study focuses only on binding affinities, which may not always perfectly reflect viral escape from antibody neutralization^31,32^. In particular, some of the binding affinities we measure could be too weak to be physiologically relevant, and mutations may impact neutralization without impacting binding affinity significantly. However, because neutralization cannot occur in the absence of affinity, our measurements are likely to be relevant for understanding the reduced sensitivity of BA.1 to these antibodies. We also note that practical constraints limit us to studying four antibodies. This limits the generalizability of our results, particularly in light of previous structural studies which have revealed more epitopes bound by mAbs.

In spite of these limitations, our binding affinity landscapes reveal that BA.1 can escape diverse antibodies by acquiring a few large-effect mutations and can reduce affinity to others by accumulating several small-effect mutations. For the first three antibodies, one or two mutations are sufficient for total escape. However, in some cases, additional mutations can restore affinity, and in others, specific combinations of large and small-effect mutations can abrogate affinity. Thus, despite the seemingly simple landscape of antibody escape, there are alternative, more intricate pathways that can abrogate affinity. In contrast, for the S309 antibody, four mutations drive the decline in affinity yet are also involved in higher-order epistatic interactions that counteract this decline. This epistasis results in an affinity threshold, beyond which additional mutations do not reduce affinity.

Predicting the future evolution of the Omicron lineage will require determining how these affinity landscapes translate to immune evasion, how antibody affinity landscapes vary within a class or epitope group, and how mutations beyond this set may further enhance immune evasion. For example, neutralization assays with minimally mutated genotypes would confirm whether the strong-effect mutations are indeed sufficient for escape. Further, assessing affinity landscapes for additional antibodies with similar epitopes would reveal how the landscape structure varies within such a group, and whether there are general features that we can extrapolate to unmeasured sequences. Finally, integrating these combinatorial libraries with saturating mutagenesis approaches would reveal how the evolvability of this lineage changes over time, and what additional mutations – such as those in BA.2, BA.4, or BA.5 – might confer further immune escape. Looking beyond the Omicron lineage, such approaches could provide more general insight into how mutations in SARS-CoV-2 may result in host-range expansion or antigenic evolution.

## METHODS

### Yeast display plasmid, strains, and library production

We used the same library and strains as produced in Moulana et al^23^. In brief, to generate clonal yeast strains for the Wuhan Hu-1 and Omicron BA.1 variants, we cloned the corresponding RBD gblock into a pETcon vector via Gibson Assembly. We then extracted and transformed Sanger-verified plasmids into the AWY101 yeast strain (kind gift from Dr. Eric Shusta)^33^ as described in Gietz and Schiestl^34^. To produce the RBD variant library, we employed a Golden Gate combinatorial assembly strategy. We constructed full RBD sequences from five sets of dsDNA fragments of roughly equal size. Each set contains versions of the fragments that differ by the mutations included. Following bacterial transformation of this Golden Gate assembly product, we extracted and transformed the library into AWY101 yeast strain, from which we inoculated and froze a library containing obtained ~1.2 million colonies.

### High-throughput binding affinity assay (Tite-Seq)

We performed Tite-seq assay as previously described^20,23,26,27^, with two replicates for each antibody (LY-CoV016, LY-CoV555, REGN10987, and S309 [Genscript, Gene-to-Antibody service]) assay on different days, for a total of eight assays.

Briefly, we thawed yeast RBD library and the Wuhan Hu-1 and Omicron BA.1 strains by inoculating the corresponding glycerol stocks in SDCAA (6.7 g/L YNB without amino acid [VWR #90004-150], 5 g/L ammonium sulfate [Sigma-Aldrich #A4418], 2% dextrose [VWR #90000– 904], 5 g/L Bacto casamino acids [VWR #223050], 1.065 g/L MES buffer [Cayman Chemical, Ann Arbor, MI, #70310], 100 g/L ampicillin [VWR # V0339])at 30°C for 20 hr. The cultures were then induced in SGDCAA (6.7 g/L YNB without amino acid [VWR #90004-150], 5 g/L ammonium sulfate [Sigma-Aldrich #A4418], 2% galactose [Sigma-Aldrich #G0625], 0.1% dextrose [VWR #90000–904], 5 g/L Bacto casamino acids [VWR #223050], 1.065 g/L MES buffer [Cayman Chemical, Ann Arbor, MI, #70310], 100 g/L ampicillin [VWR # V0339]), and rotated at room temperature for 16–20 hr.

Following overnight induction, we pelleted, washed (with 0.01% PBSA [VWR #45001–130; GoldBio, St. Louis, MO, #A-420–50]), and incubated the cultures with monoclonal antibody at a range of concentrations (10^−6^ to 10^−12^ with 0.75-log increments for CoV555, 10^−7^ to 10^−12^ with 0.5-log increments for S309, 10^−6^ to 10^−12.7^ with 0.75-log increments for REGN10987, 10^−6^ to 10^−12^ with 0.75-log increments for SB6). The yeast-antibody mixtures were incubated at room temperature for 20 hr. The cultures were then pelleted washed twice with PBSA and subsequently labeled with PE-conjugated goat anti-human IgG (1:100, Jackson ImmunoResearch Labs #109-115-098) and FITC-conjugated chicken anti-cMmyc (1:100, Immunology Consultants Laboratory Inc., Portland, OR, #CMYC-45F). The mixtures were rotated at 4°C for 45 min and then washed twice in 0.01% PBSA.

Sorting, recovery, and sequencing library preparation followed Moulana, et al^23^. In short, we sorted ~1.2 million yeast cells per concentration, gated by FSC vs SSC and then by expression (FITC) and/or binding fluorescence (PE) on a BD FACS Aria Illu. The machine was equipped with 405 nm, 440 nm, 488 nm, 561 nm, and 635 nm lasers, and an 85 micron fixed nozzle. Sorted cells were then pelleted, resuspended in SDCAA, and rotated at 30°C until late-log phase (OD600 = 0.9-1.4). The cultures were then pelleted and stored at −20°C for at least six hours prior to extraction using Zymo Yeast Plasmid Miniprep II (Zymo Research # D2004), following the manufacturer’s protocol. The sequencing amplicon libraries were then prepared by a two-step PCR as previously described^23,27,35^. In brief, we added to the amplicon unique molecular identifies (UMI), inline indices, and partial Illumina adapters through a 7-cycle PCR which amplifies the RBD sequence in the plasmid. We then used the cleaned product from the first PCR in the second PCR to append Illumina i5 and i7 indices accordingly (see https://github.com/desai-lab/compensatory_epistasis_omicron/tree/main/Supplementary_Files for primer sequences). The products were then cleaned using 0.85x Aline beads, verified using 1% agarose gel, quantified on Spectramax i3, pooled, and verified on Tapestation 5000HS and 1000HS. Final library was quantitated by Qubit fluorometer and sequenced on Illumina Novaseq SP, supplemented with 10% PhiX.

### Sequence data processing

Following Moulana et al.^23^, we processed raw demultiplexed sequencing reads to identify and extract the indexes and mutational sites. Briefly, for each antibody, we utilized a snakemake pipeline (https://github.com/desai-lab/omicron_ab_landscape) to parse through all fastq files and group the reads according to inline indices, UMIs, and sequence reads. We accepted sequences based on criteria previously determined (10% bp mismatches) and converted accepted sequences into binary genotypes (‘0’ for Wuhan Hu-1 allele or ‘1’ for Omicon BA.1 allele at each mutation position). Reads containing errors at mutation sites were removed. Finally, the pipeline collated genotype counts based on distinct UMIs from all samples into a single table.

We fit the binding dissociation constants *K*_D,app_ for each genotype as previously described^23,27^. Briefly, using sequencing and flow cytometry data, we calculated the mean log-fluorescence of each genotype *s* at each concentration *c*, as follows:

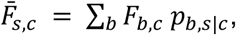

where *F_b,c_* is the mean log-fluorescence of bin *b* at concentration *c*, and *p_b,s|c_* is the inferred proportion of cells from genotype *s* that are sorted into bin *b* at concentration *c*, which is estimated from the read counts as:

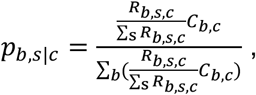

Here, *R_b,s,c_* represents the number of reads from genotype *s* that are found in bin *b* at concentration *c*, and *C_b,c_* refers to the number of cells sorted into bin *b* at concentration *c*.

We then computed the uncertainty for the mean log-fluorescence:

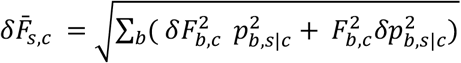

where *δF_b,c_* is the spread of the log fluorescence of cells sorted into bin *b* at concentration *c*. The error in *p_b,s|c_* emerges from the sampling error, which can be approximated as a Poisson process, such that:

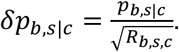

Finally, we inferred the binding dissociation constant (*K*_D,s_) for each variant by fitting the logarithm of Hill function to the mean log-fluorescence 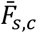, as a function of concentrations *c*:

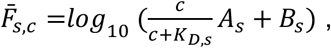

where *A*_s_ is the increase in fluorescence at antibody saturation, and *B*_s_ is the background fluorescence level. The fit was performed using the *curve_fit* function in the Python package *scipy.optimize*. Across all genotypes, we imposed bounds on the values of *A*_s_ to be 10^2^-10^6^, *B*_s_ to be 1-10^5^, and *K*_D,s_ to be 10^−14^-10^−5^. We then averaged the inferred *K*_D,s_ values across the two replicates for each antibody after removing values with poor fit (*r*^2^ < 0.8 or standard error > 1). Variants were defined as non-binders if the difference between the maximum and the minimum of their estimated log-fluorescence over all concentrations was lower than 1 (in log-fluorescence units). This value was set by measuring the distribution for known non-binders (see Supplementary Figure S3).

### Isogenic measurements for validation

We validated our high-throughput binding affinity method by measuring the binding affinities for the Wuhan Hu-1 and Omicron BA.1 RBD variants. For each isogenic titration curve, we followed the same labeling strategy as in Tite-seq, titrating each antibody at concentrations ranging from 10^−12^-10^−7^ M (with increments of 0.5 for the first replicate and 1 for the second one) for isogenic yeast strains that display only the sequence of interest. The mean log fluorescence was measured using a BD LSR Fortessa cell analyzer. We directly computed the mean and variances of these distributions for each concentration and used them to infer the value of *K*_D,app_ using the formula shown above.

### Decision trees on loss-of-binding mutations

To summarize mutations that drive the loss of binding (escape) for each antibody, we constructed a decision tree using package rpart in R^36^ with its default parameters. In brief, for each antibody (except for S309 where every sequence binds the antibody), we first categorized each genotype into a binary parameter with values ‘binding’ or ‘non-binding’. Then, the function rpart splits the tree based on any one of the fifteen mutations by minimizing the Gini impurity for the binding parameter. The method continues to partition the tree if the cost complexity parameter (cp) of the split does not drop below 0.01. This parameter is the sum of all misclassifications (binding vs. non-binding) at every terminal node (analogous to residual sum of squares in regression), added by the product between the number of splits (analogous to degree of freedom) and a penalty term inferred through cross-validation performed by the rpart algorithm. The tree is then presented in Figure 2 using `ggparty` package^37^.

### Epistasis analysis

We used a linear model where the effects of combinations of mutations sum to the phenotype of a sequence. The logarithm of the binding affinity log *K*_D,app_ is proportional to change in free energy. Thus, without epistatic interactions, the effects of mutations are expected to combine additively^38,39^. However, some phenotypic values log *K*_D,app_ are unavailable in our dataset due to the upper limit of the assay concentration. We are unable to precisely determine *K*_D,app_ for the low-affinity (or non-binding) variants, especially when the true −log *K*_D,app_ < 6.0 (the highest log-concentration used). To deal with this problem, we supplemented our linear model with a boundary, following tobit left-censored model. In this model, the −log *K*_D,app_ phenotype is measured as 6.0 for all values lower than 6.0. Thus, the full *K*-order model can be written as:

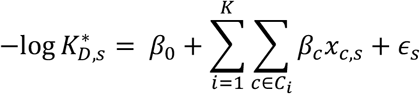

where *C*_*i*_ contains all 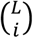 combinations of size *i* of the mutations and *x*_c,s_ equal to 1 if the sequence *s* contains all the mutations in *c* and to 0 otherwise. Here, 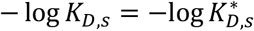 if 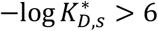 and − log *K*_*D,s*_ = 6 if 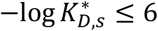. Then, following the tobit model approach, we compute the likelihood function to infer coefficient parameters 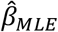, given by:

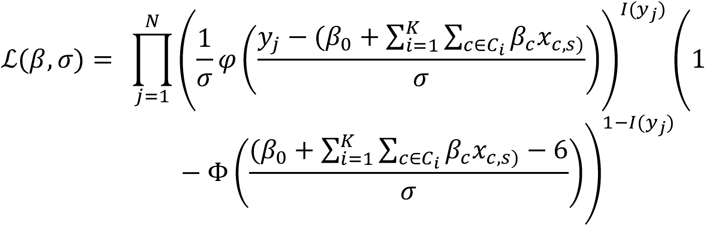

where *y*_*j*_ = − log *K*_D,app,*j*_, and *ϕ* and Φ denote the standard normal cumulative distribution function and probability density function, respectively. Moreover, note that 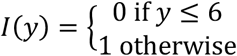. This optimization problem would include coefficients that are associated with the loss-of-binding phenotypes. Consequently, by the model, these coefficients do not have lower bounds and the optimization would have resulted in deflated coefficients offset by inflated higher-order coefficients, or vice-versa. To resolve this problem, we add a lasso regularization term in the form of *ϵ* ∑|*β*_*c*_| to the likelihood, with *ϵ* = 0.01. This term is small enough to reduce the magnitude of constrained coefficients but act as intended on the non-constrained ones. To maximize the log-likelihood function, which is a concave function, we used the optimize module in the scipy package, with the BFGS gradient-descent method.

### Structural analysis

We used the reference structure of a 2.79 Å cryo-EM structure of Omicron BA.1 complexed with (PDB ID: 7WPB). The contact surface area is determined by using ChimeraX^40^ to measure the buried surface area between ACE2 and each mutated residue in the RBD (*measure buriedarea* function, default probeRadius of 1.4Å), whereas distance between α-carbons is measured using PyMol^41^.

### Force directed layout

The high-dimensional binding affinity landscape can be projected in two dimensions with a force-directed graph layout approach (see https://desai-lab.github.io/wuhan_to_omicron/). Each node corresponds to each sequence in the library, connected by edges to a neighbor that differs in one single site. For each antibody, an edge between two sequences *s* and *t* is given the weight:

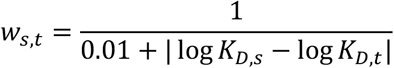

Additionally, we also constructed a different layout that includes affinities to all antibodies, where the weight between two sequences depends on the sum over the antibodies of the difference between their affinities:

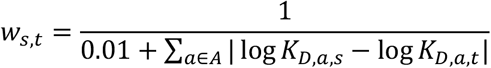

where *A* is the set of antibodies we used. In a force-directed representation, the edges pull together the nodes they are attached to proportional to the weight given to each edge. In our scenario, this means that nodes with a similar genotype (a few mutations apart) and a similar phenotype (binding affinity or total binding affinity) will be close to each other in two dimensions.

Importantly this is not a “landscape” representation: the distance between two points is unrelated to how easy it is to reach one genotype from another in a particular selection model.

Practically, after assigning all edge weights, we use the layout function *layout_drl* from the Python package *iGraph*, with default settings, to obtain the layout coordinates for each variant.

### Genomic data

To analyze SARS-CoV-2 phylogeny, we used all complete RBD sequences from all SARS-CoV-2 genomes deposited in the Global Initiative on Sharing All Influenza Data (GISAID) repository^42–44^ with the GISAID Audacity global phylogeny (EPI_SET ID: EPI_SET_20220615uq, available on GISAID up to June 15, 2022, and accessible at https://doi.org/10.55876/gis8.220615uq). We pruned the tree to remove all sequences with RBD not matching any of the possible intermediates between Wuhan Hu-1 and Omicron BA.1 and analyzed this tree using the python toolkit ete3^45^. We measured the frequency of each mutation by counting how many times it emerges in the tree, normalized by the total occurrences of other mutations. For frequency with Q498R and N501Y, we counted the occurrence of each mutation only on branches that already contains Q498R and N501Y and normalized similarly.

### Statistical analyses and visualization

All data processing and statistical analyses were performed using R v4.1.0^46^ and python 3.10.047. All figures were generated using ggplot2^48^ and matplotlib^49^.

## ACKNOWLEDGEMENTS

We thank Zach Niziolek for flow cytometry assistance and all members of the Desai lab and Serafina Nieves for helpful discussions. T.D. acknowledges support from the Human Frontier Science Program Postdoctoral Fellowship, A.M.P. acknowledges support from the Howard Hughes Medical Institute Hanna H. Gray Postdoctoral Fellowship, J.C. acknowledges support from the National Science Foundation Graduate Research Fellowship, and M.M.D. acknowledges support from the NSF-Simons Center for Mathematical and Statistical Analysis of Biology at Harvard University, supported by NSF grant no. DMS-1764269, and the Harvard FAS Quantitative Biology Initiative, grant DEB-1655960 from the NSF and grant GM104239 from the NIH. J.D.B. acknowledges support from NIH/NIAID grant R01AI141707 and is an Investigator of the Howard Hughes Medical Institute. We gratefully acknowledge all data contributors, i.e. the Authors and their Originating laboratories responsible for obtaining the specimens, and their Submitting laboratories for generating the genetic sequence and metadata and sharing via the GISAID Initiative. Computational work was performed on the FASRC Cannon cluster supported by the FAS Division of Science Research Computing Group at Harvard University.

## AUTHOR CONTRIBUTIONS

Conceptualization: A.M., T.D., A.M.P., J.C., T.N.S., A.J.G., J.D.B., and M.M.D. Methodology: A.M., T.D., A.M.P., J.C., S.N., T.N.S., and A.J.G. Library design and production: A.M., T.D., A.M.P., J.C., and A.J.G. Experiments: A.M., T.D., A.M.P., J.C., and A.A.R. Validation: A.M., T.D., A.M.P., J.C., S.N., and T.N.S. Data analysis: A.M., T.D., A.M.P., J.C., S.N., and T.N.S. Supervision: A.M.P, J.D.B., and M.M.D. Funding acquisition: J.D.B. and M.M.D. Writing— original draft: A.M., T.D., A.M.P., J.C., and M.M.D. All the authors reviewed and edited the manuscript.

## DATA AND CODE AVAILABILITY STATEMENT

Raw sequencing reads have been deposited in the NCBI BioProject database under accession number PRJNA877045. All associated metadata are available at https://github.com/desai-lab/omicron_ab_landscape.

---

**Supplementary Figure S1.**
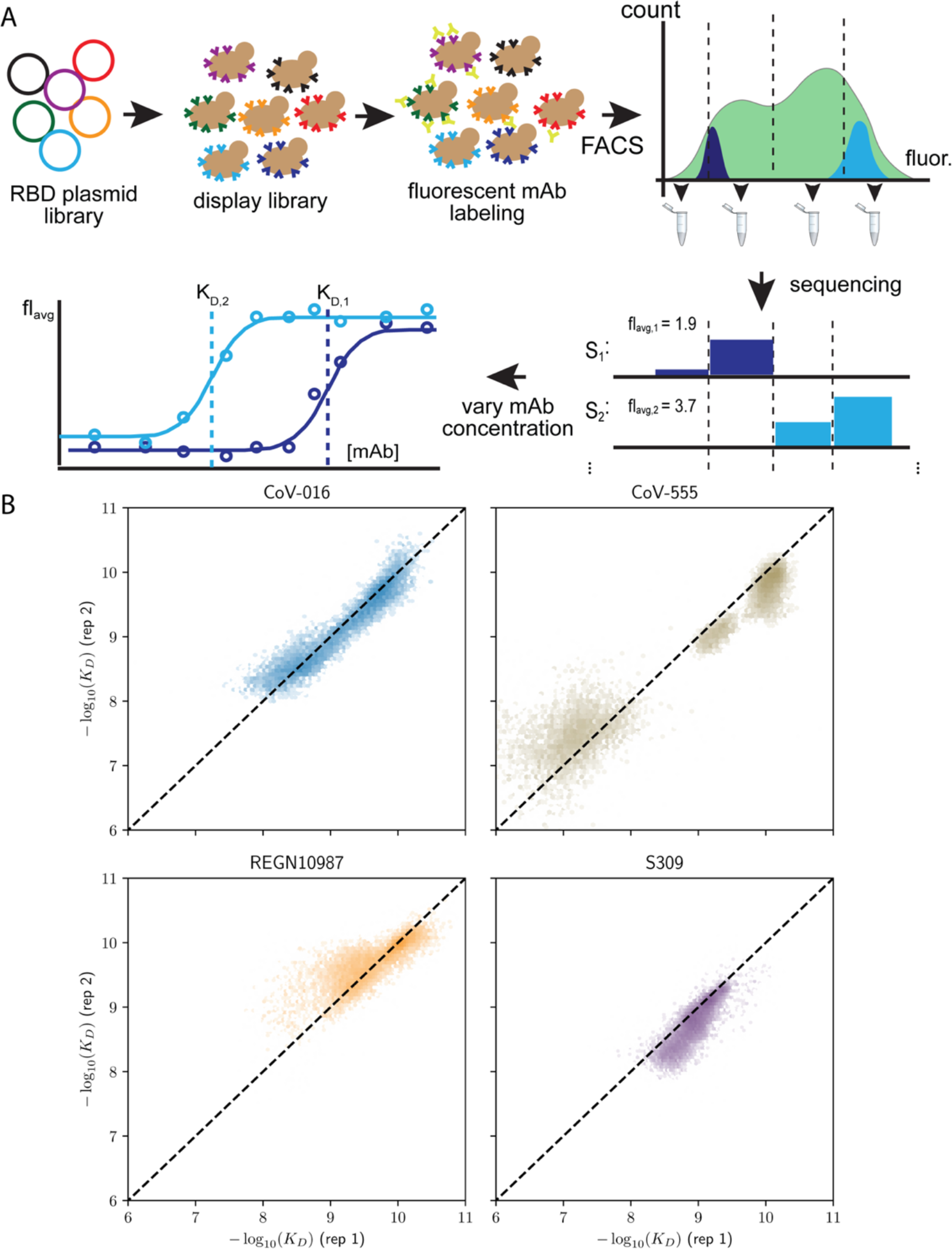
Schematic overview of the Tite-Seq method and reproducibility of dissociation constants. (**A**) The plasmid library containing RBD sequences is transformed into a standard yeast display strain AWY101. The library is then incubated with soluble, fluorescent monoclonal antibody (either one of the four mAbs used in the study) and sorted by flow cytometry into bins based on mAb fluorescence. Deep sequencing of each bin results in the mean bin (B_avg_) estimate of each RBD variant across varying mAb concentrations to produce a titration curve. Apparent equilibrium dissociation constants are them inferred by fitting B_avg_ to the mAb concentration. (**B**) Correlation of −log(*K*_D,app_) between the first and second biological replicates.

**Supplementary Figure S2.**
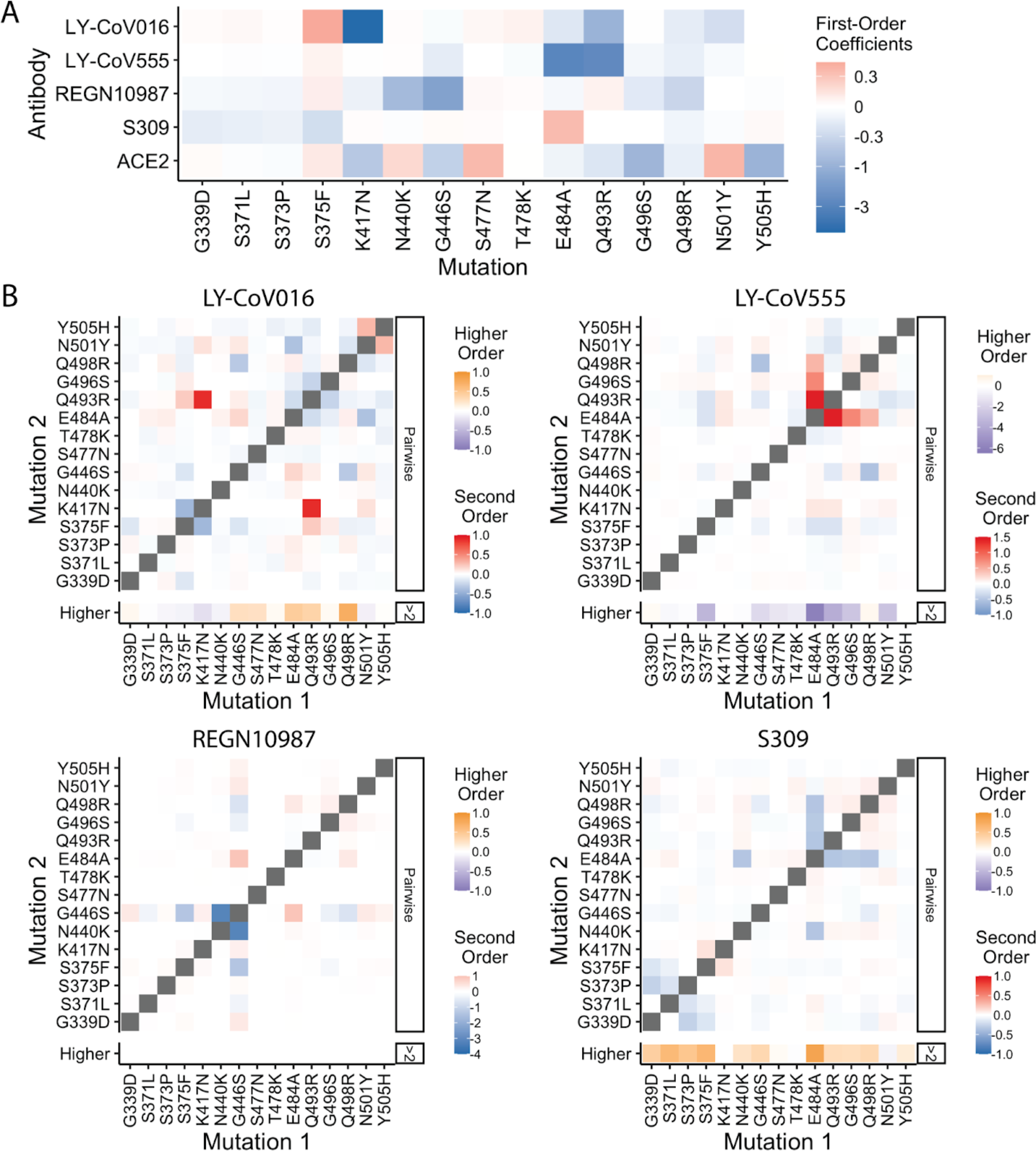
Epistatic coefficients. (**A**) First-order effects in fourth order epistatic interaction models. (**B**) For each antibody, higher-order interaction coefficient of a mutation (shown at bottom of heat map plot) is calculated by summing over all third- and fourth-order interaction coefficients involving the mutation.

**Supplementary Figure S3.**
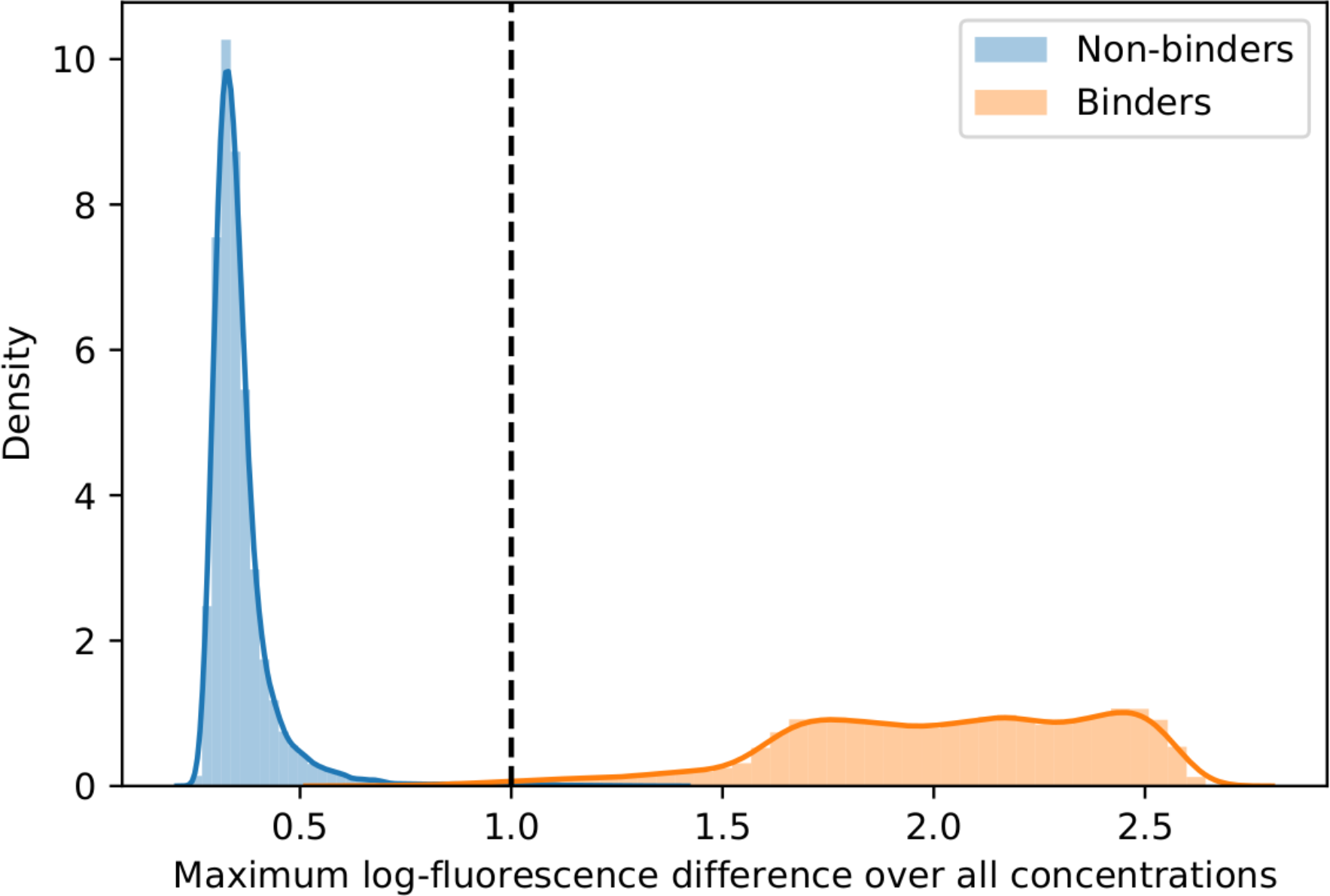
Distribution of maximum log-fluorescence difference. To determine whether a genotype is a non-binder, the maximum difference of log fluorescence across concentrations of the genotype is computed. The dashed line represents the threshold below which a genotype with a certain maximum difference is considered non-binding.

**Supplementary Table S1.** Isogenic validation of binding affinities. The *K*_D,app_ inferred from Isogenic measurements (see Methods) shown with those inferred via Tite-Seq measurement. NB denotes non-binding and standard deviations between replicates are also shown.

| Strain | Antibody | Isogenic -log<br>$K_{D,app}$ | TiteSeq -log<br>$K_{D,app}$ |
| --- | --- | --- | --- |
| Omicron BA.1 | LY-CoV016 | NB | NB |
| Omicron BA.1 | LY-CoV555 | NB | NB |
| Omicron BA.1 | REGN10987 | NB | NB |
| Omicron BA.1 | S309 | 8.81 | $8.59 \pm 0.16$ |
| Wuhan Hu-1 | LY-CoV016 | $10.52 \pm 0.24$ | $9.80 \pm 0.02$ |
| Wuhan Hu-1 | LY-CoV555 | $10.01 \pm 0.33$ | $10.19 \pm 0.24$ |
| Wuhan Hu-1 | REGN10987 | $10.42 \pm 0.49$ | $9.96 \pm 0.05$ |
| Wuhan Hu-1 | S309 | $9.33 \pm 0.27$ | $9.34 \pm 0.09$ |

## REFERENCES CITED

1. Cao, Y. et al. Omicron escapes the majority of existing SARS-CoV-2 neutralizing antibodies. Nature 602, 657–663 (2022).

2. Ao, D. et al. SARS-CoV-2 Omicron variant: Immune escape and vaccine development. MedComm (2020) 3, e126 (2022).

3. Planas, D. et al. Considerable escape of SARS-CoV-2 Omicron to antibody neutralization. Nature 602, 671–675 (2022).

4. Viana, R. et al. Rapid epidemic expansion of the SARS-CoV-2 Omicron variant in southern Africa. Nature 603, 679–686 (2022).

5. Greaney, A. J. et al. Mapping mutations to the SARS-CoV-2 RBD that escape binding by different classes of antibodies. Nat. Commun. 12, 4196 (2021).

6. Greaney, A. J. et al. Complete mapping of mutations to the SARS-CoV-2 spike receptor-binding domain that escape antibody recognition. Cell Host Microbe 29, 44–57.e9 (2021).

7. Iketani, S. et al. Antibody evasion properties of SARS-CoV-2 Omicron sublineages. Nature 604, 553–556 (2022).

8. Dai, L. & Gao, G. F. Viral targets for vaccines against COVID-19. Nat. Rev. Immunol. 21, 73–82 (2021).

9. Barnes, C. O. et al. Structures of human antibodies bound to SARS-CoV-2 spike reveal common epitopes and recurrent features of antibodies. Cell 182, 828–842.e16 (2020).

10. Barnes, C. O. et al. SARS-CoV-2 neutralizing antibody structures inform therapeutic strategies. Nature 588, 682–687 (2020).

11. Liu, C. et al. Reduced neutralization of SARS-CoV-2 B.1.617 by vaccine and convalescent serum. Cell 184, 4220–4236.e13 (2021).

12. Zhou, D. et al. Evidence of escape of SARS-CoV-2 variant B.1.351 from natural and vaccine-induced sera. Cell 184, 2348–2361.e6 (2021).

13. Greaney, A. J. et al. The SARS-CoV-2 Delta variant induces an antibody response largely focused on class 1 and 2 antibody epitopes. PLoS Pathog. 18, e1010592 (2022).

14. Greaney, A. J. et al. Comprehensive mapping of mutations in the SARS-CoV-2 receptor-binding domain that affect recognition by polyclonal human plasma antibodies. Cell Host Microbe 29, 463–476.e6 (2021).

15. Dejnirattisai, W. et al. SARS-CoV-2 Omicron-B.1.1.529 leads to widespread escape from neutralizing antibody responses. Cell 185, 467–484.e15 (2022).

16. Cameroni, E. et al. Broadly neutralizing antibodies overcome SARS-CoV-2 Omicron antigenic shift. Nature 602, 664–670 (2022).

17. Starr, T. N. et al. Prospective mapping of viral mutations that escape antibodies used to treat COVID-19. Science 371, 850–854 (2021).

18. Starr, T. N. et al. SARS-CoV-2 RBD antibodies that maximize breadth and resistance to escape. Nature 597, 97–102 (2021).

19. Chakraborty, C., Sharma, A. R., Bhattacharya, M. & Lee, S.-S. A detailed overview of immune escape, antibody escape, partial vaccine escape of SARS-CoV-2 and their emerging variants with escape mutations. Front. Immunol. 13, 801522 (2022).

20. Starr, T. N. et al. Deep mutational scanning of SARS-CoV-2 receptor binding domain reveals constraints on folding and ACE2 binding. Cell 182, 1295–1310.e20 (2020).

21. Mannar, D. et al. SARS-CoV-2 Omicron variant: Antibody evasion and cryo-EM structure of spike protein-ACE2 complex. Science 375, 760–764 (2022).

22. McCallum, M. et al. Structural basis of SARS-CoV-2 Omicron immune evasion and receptor engagement. Science 375, 864–868 (2022).

23. Moulana, A. et al. Compensatory epistasis maintains ACE2 affinity in SARS-CoV-2 Omicron BA.1. bioRxiv (2022) doi:10.1101/2022.06.17.496635.

24. Starr, T. N. et al. Shifting mutational constraints in the SARS-CoV-2 receptor-binding domain during viral evolution. Science 377, 420–424 (2022).

25. Case, J. B. et al. Resilience of S309 and AZD7442 monoclonal antibody treatments against infection by SARS-CoV-2 Omicron lineage strains. Nat. Commun. 13, 3824 (2022).

26. Adams, R. M., Mora, T., Walczak, A. M. & Kinney, J. B. Measuring the sequence-affinity landscape of antibodies with massively parallel titration curves. Elife 5, (2016).

27. Phillips, A. M. et al. Binding affinity landscapes constrain the evolution of broadly neutralizing anti-influenza antibodies. Elife 10, (2021).

28. Starr, T. N. et al. ACE2 binding is an ancestral and evolvable trait of sarbecoviruses. Nature 603, 913–918 (2022).

29. Windsor, I. W. et al. Antibodies induced by an ancestral SARS-CoV-2 strain that cross-neutralize variants from Alpha to Omicron BA.1. Sci. Immunol. 7, eabo3425 (2022).

30. Javanmardi, K. et al. Antibody escape and cryptic cross-domain stabilization in the SARS-CoV-2 Omicron spike protein. Cell Host Microbe (2022) doi:10.1016/j.chom.2022.07.016.

31. Steckbeck, J. D. et al. Kinetic rates of antibody binding correlate with neutralization sensitivity of variant simian immunodeficiency virus strains. J. Virol. 79, 12311–12320 (2005).

32. Culp, T. D., Spatz, C. M., Reed, C. A. & Christensen, N. D. Binding and neutralization efficiencies of monoclonal antibodies, Fab fragments, and scFv specific for L1 epitopes on the capsid of infectious HPV particles. Virology 361, 435–446 (2007).

33. Wentz, A. E. & Shusta, E. V. A novel high-throughput screen reveals yeast genes that increase secretion of heterologous proteins. Appl. Environ. Microbiol. 73, 1189–1198 (2007).

34. Gietz, R. D. & Schiestl, R. H. Quick and easy yeast transformation using the LiAc/SS carrier DNA/PEG method. Nat. Protoc. 2, 35–37 (2007).

35. Nguyen Ba, A. N. et al. High-resolution lineage tracking reveals travelling wave of adaptation in laboratory yeast. Nature 575, 494–499 (2019).

36. Therneau, T., Atkinson, B., & Ripley, B. Rpart: Recursive Partitioning. (2013).

37. Borkovec, M. et al. ggparty: “ggplot” Visualizations for the “partykit” Package. (2019).

38. Wells, J. A. Additivity of mutational effects in proteins. Biochemistry 29, 8509–8517 (1990).

39. Olson, C. A., Wu, N. C. & Sun, R. A comprehensive biophysical description of pairwise epistasis throughout an entire protein domain. Curr. Biol. 24, 2643–2651 (2014).

40. Pettersen, E. F. et al. UCSF ChimeraX: Structure visualization for researchers, educators, and developers. Protein Sci. 30, 70–82 (2021).

41. Schrodinger, L. L. C. The PyMOL Molecular Graphics System. (2015).

42. Khare, S. et al. GISAID’s role in pandemic response. China CDC Wkly 3, 1049–1051 (2021).

43. Elbe, S. & Buckland-Merrett, G. Data, disease and diplomacy: GISAID’s innovative contribution to global health. Global Chall. 1, 33–46 (2017).

44. Shu, Y. & McCauley, J. GISAID: Global initiative on sharing all influenza data – from vision to reality. Euro Surveill. 22, (2017).

45. Huerta-Cepas, J., Serra, F. & Bork, P. ETE 3: Reconstruction, analysis, and visualization of phylogenomic data. Mol. Biol. Evol. 33, 1635–1638 (2016).

46. R Core Team. R: A language and environment for statistical computing. (2017).

47. Van Rossum, G. & Drake, F. L. Python 3 Reference Manual. (CreateSpace, 2009).

48. Wickham, H. Ggplot2. (Springer International Publishing, 2016).

49. Hunter, J. D. Matplotlib: A 2D graphics environment. Computing in Science & Engineering 9, 90–95 (2007).

